# Engineering gene drive docking sites in a haplolethal locus in *Anopheles gambiae*

**DOI:** 10.1101/2025.03.03.641265

**Authors:** Andrea L. Smidler, Eryney A. Marrogi, Sean Scott, Enzo Mameli, Daniel Abernathy, Omar S. Akbari, George M. Church, Flaminia Catteruccia, Kevin Esvelt

## Abstract

Gene drives are selfish genetic elements which promise to be powerful tools in the fight against vector-borne diseases such as malaria. We previously proposed population replacement gene drives designed to better withstand the evolution of resistance by homing through haplolethal loci. Because most mutations in the wild-type allele that would otherwise confer resistance are lethal, only successful drive homing permits the cell to survive. Here we outline the development and characterization of two ΦC31-Recombination mediated cassette exchange (RMCE) gene drive docking lines with these features in *Anopheles gambiae*, a first step towards construction of robust gene drives in this important malaria vector. We outline adaption of the technique HACK (Homology Assisted CRISPR knockin) to knock-in two docking site sequences into a paired haplolethal-haplosufficient (Ribosome-Proteasome) locus, and confirm that these docking lines permit insertion of drive-relevant transgenes. We report the first anopheline proteasome knockouts, and identify ribosome mutants that reveal a major hurdle that such designs must overcome to develop robust drives in the future. Although we do not achieve drive, this work provides a new tool for constructing future evolution-robust drive systems and reveals critical challenges that must be overcome for future development of gene drives designed to target haplolethal loci in anophelines and, potentially, other metazoans.

## BACKGROUND

Gene drive technology could revolutionize the control of vector-borne diseases [1,2]. Designed to either suppress wild populations or replace them with disease-refractory ones, gene drives have been proposed as a viable tool to control diseases such as malaria [3]. Homing gene drives (HGD) are artificial selfish genetic elements capable of copying themselves from one homolog to the other by homology directed repair (HDR). This copying is catalyzed by targeted CRISPR cleavage of the non-drive chromosome and repair of the break with the drive transgenic cassette. If this occurs in the germline, this results in Super-Mendelian inheritance of the gene drive cassette, spreading desirable genetic cargoes or traits throughout the population. HGDs can therefore be used to spread sterilizing or antipathogenic traits, resulting in population suppression or population replacement respectively, to achieve different vector control outcomes.

Inherent to any HGD design is the generation of resistance alleles at the target locus. Generated as a byproduct of end-joining repair following CRISPR cleavage, they are an inevitable hindrance of many drive designs [4–7]. Resistant-conferring polymorphisms prevent subsequent guide RNA (gRNA) binding and cleavage, preclude future drive, and often compete the drive to extinction [6–9]. Though some drives target haplosufficient genes to provide some selection against resistance allele formation [10,11], all gene drives developed in anophelines to date have induced some level of resistance [3–21]. Resistance alleles can be categorized into two types; r*1* resistance alleles are those which preserve the function of the target gene, while *r2* alleles are those which disrupt gene function. Both must be prevented to guarantee drive spread [22]. Our previously proposed drive designs, termed Evolutionarily Stable Homing Gene Drives (ESHGDs), use multiple gRNAs to home into a haplolethal target. In doing so they can theoretically prevent both types of resistance [1,23,24], as r1 alleles are deleted during homing and r2 alleles are lethal. This design causes only correct drive homing to be survivable for the cell. ESHGDs are not to be confused with other drive designs targeting a haplolethal locus, whose long-term success is unlikely. In these designs there are additional homologous sequences within the recoded region, making HDR with these sequences likely, causing transgene shuffling[25] and probable failure of this drive design.

Recombinase Mediated Cassette Exchange (RMCE) is a technology for docking desired cargoes onto ΦC31 attB or attP sequences, enabling precise insertion of genetic cargoes onto target sequences. Here we report on the generation of two gene drive RMCE docking lines with the recoding and genetic engineering necessary to enable the creation of ESHGDs. We set out to generate docking lines within the dual-gene haplolethal-haplosufficient gene locus, Ribosome Protein L11 and Proteasome Regulatory Particle Subunit 1 (*RpL11-Rpt1)*. In doing so we replace the 3’-terminal coding and UTR sequences with recoding and replacement UTRs, while providing ΦC31 RMCE docking sequences for subsequent transgenesis of drives. We demonstrate that one of these docking lines is an expected proteasome knockout, while the other is an unexpected Ribosome Minute mutant. While we are unable to demonstrate homing, we demonstrate that these docking lines can support transgenesis of a drive-like transgene, and may be able to support a drive following future fine-tuning of Cas9 expression. In all, this work presents an important advancement in the development of gene drives that are less susceptible to resistance.

## RESULTS

### Design of two gene drive docking sites, dRP and dRPi

Our ESHGD designs target haplolethal genes to make any non-drive outcome lethal for the cell. In these designs the 3’-terminal coding sequence and 3’UTR are deleted from a haplolethal gene and are replaced with recoded coding sequence and a 3’UTR from a similarly regulated paralog. This design is meant to reconstitute gene function while creating a region not shared with the wild type homolog, which can be uniquely targeted with multiple gRNAs, over which HDR can not occur (**Figure 1A, B**). In a subsequent transgenesis step, CRISPR-encoding drive sequences are inserted immediately adjacent to the recoded haplolethal gene, without disrupting it. By targeting the wild type haplolethal coding sequence, such drives are intolerant of *r2* mutations due to the essentiality of the haplolethal gene product [4], and simultaneously delete any *r1* alleles which do arise during homing due to the HDR-inhibition caused by the recoding. This design theoretically circumvents the issue of resistance which has plagued most drives to date [4,9,22,26–28], exhibiting a true delete-and-paste function [29] in which only successful drive homing allows the cell to survive. This makes inheritance of the recoded gene ‘addictive’[30].

**Figure 1.**
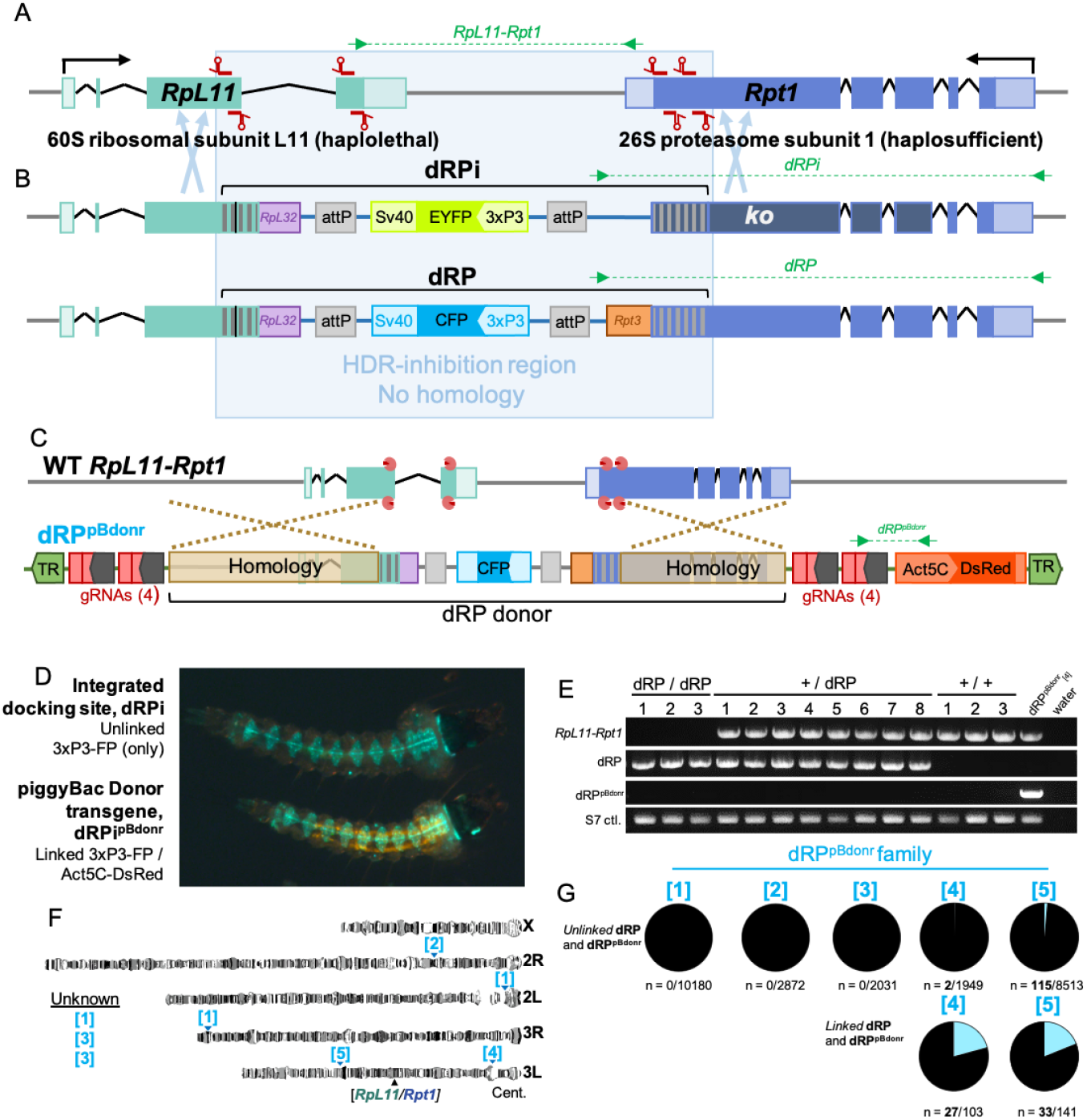
Design and HACK-based integration of dRPi and dRP into *RpL11-Rpt1*. **[A]** The endogenous wild type *RpL11-Rpt1* target locus on Chromosome 3L shown to scale. Location of 8 gRNA binding sites (red) targeting the C-terminal coding sequence of each gene. **[B]** Design of dRPi and dRP docking sites, not to scale. Codon recoding (vertical grey bars) corresponds to the C-terminal 183 bp and 258 bp of *RpL11* and *Rpt1* respectively. *RpL11* intron 3 was removed (vertical black bar), and replacement surrogate 3’UTR from *RpL32* provided (209 bp, purple). In dRP a replacement 3’UTR from *Rpt3* is provided to *Rpt1* to reconstitute gene function (orange), in dRPi it is omitted to eliminate *Rpt1* function. Also included are attP sequences for ΦC31 transgenesis of subsequent drive designs by RCME (grey), and fluorescent selectable markers, *3xP3-EYFP* or *3xP3-CFP (m2Turquoise)*, distinguish dRPi and dRP respectively. HDR-inhibition region is outlined (pale blue). Pale blue arrows show region where HDR can occur. **[C]** Design of dRP^pBdonr^ piggyBac donor transgene for HACK integration of dPR into *RpL11-Rpt1*, and scheme of knock-in. The donor region (black bracket) encompasses 2,241bp and 1,550bp hompology arms for *RpL11* and *Rpt1* respectively (beige) and dRPi and dRP docking sequences (dRP shown). Outside the donor includes 8 gRNAs as 4 tethered pairs (red) under expression of the PolII promoter (dark grey arrows). CRISPR targets WT RpL11-Rpt1 shown with Cas9 (pink Pac-man) and gRNAs (red). **[D]** Act5c-DsRed/3xP3-FP fluorescence is indicative of the donor transgene. HACK-based integration yields larvae with 3xP3-FP fluorescence alone. **[E]** PCR validation for dRP insertion, and verification of homozygous, heterozygous and wild type adult siblings present in a mixed population cage. Three, 7-day old homozygous adults adjacent to heterozygous and wild type siblings shown. *WT RpL11-Rpt1*; primers specific to unintegrated endogenous locus. *dRP*; primers specific to dRP inserted within *RpL11-Rpt1. dRPu*^*pBdonr*^ ; primers specific to piggyBac donor transgene, spanning *Act5C-DsRed. S7*; positive control for DNA, primers specific to ribosome S7 sequence. **[F]** Individual genomic insertion sites of dRP^pBdonr^ in five families on graphical representation of *An. gambiae* polytene chromosomes. The dRP^pBdonr^ _[1]_, dRP^pBdonr^ _[2]_, dRP^pBdonr^_[3]_, dRP^pBdonr^_[4],_ and dRP^pBdonr^_[5]_ family names shortened here to [1], [2], [3], [4], and [5], respectively for brevity. Lines with insertion site sequences corresponding to unknown loci in the annotated genome summarized under *Unknown*. Endogenous *RpL11-Rpt1* locus on chromosome 3L marked for reference. **[G]** *Unlinked dRP and dRP*^*pBdonr*^ : Frequency of larvae with *3xP3-CFP* fluorescence visibly unlinked from *Act5C-DsRed* among offspring of males undergoing HACK in the germline (ie. male genotype *{*dRP^pBdonr^ /VasCas}). *Linked dRP and dRP*^*pBdonr*^ : The frequency of linked HACK events among those families which demonstrated fluorescently visible HACK (dRP^pBdonr^ _[4],_ and dRP^pBdonr^_[5]_). PCR for dRP HACK-based gene conversion insertion within larvae which co-inherited dRP^pBdonr^. visible HACK (dRP^pBdonr^ _[4],_ and dRP^pBdonr^_[5]_). PCR for dRP HACK-based gene conversion insertion within larvae which co-inherited dRP^pBdonr^.

Optimal ESHGD designs span two adjacent haplolethal genes positioned in a 3’-to-3’ orientation to create distinct recoded boundaries for homing on each side of the drive cassette. While Ribosomal subunit proteins [31], due to the need for 10 million copies per cell [32], effectively guarantee homolog haplolethality, a 3’-to-3’ Ribosome-Ribosome gene pair does not exist in the *A. gambiae* genome. We therefore selected the Ribosome-Proteasome gene pair *RpL11-Rpt1* as the target for our evolutionarily stable gene drive docking lines (**Figure 1A**). *RpL11*(AGAP011173) is a 60S well-characterized ribosomal subunit [31], whose stop codon is only 1128 bp from the stop codon of *Rpt1*, a likely haplosufficient essential ortholog 26S proteasome subunit T1 [33], making these genes a good candidate pair. Due to the complexity of the genome engineering required, we originally set out to design the gene drive in two steps. In the first step the gene drive docking lines would carry the necessary recoding, engineering, and RMCE-compatible ΦC31 docking sequences (reported herein) [3], prior to constructing a full gene drive in the locus in a later second step.

In selecting the *RpL11-Rpt1* gene pair, we developed two docking site designs. The first design sought to maintain endogenous function of both *RpL11* and *Rpt1* to facilitate ‘classic’ population replacement gene drives characterized by homozygous viability. For this we recoded 183 bp and 253 bp of *RpL11* and *Rpt1* respectively [34], removed intron 3 from *RpL11*, and provided surrogate 3’UTRs from *RpL32* and *Rpt3* respectively (**Table S1**). Further included are two attP ΦC31 sites for RMCE, and a 3xP3-CFP marker for selection. We named this docking line **<u>dRP</u>** (**d**ocking site in **<u>R</u>**ibosome and **<u>P</u>**roteasome) (**Figure 1B**). However, because the *Rpt1* homolog is likely haplosufficient, heterozygous *r2* alleles could persist in driving populations with this docking site design. We therefore designed a second docking site with *Rpt1* knocked-out in the drive-containing chromosome to render the remaining wild type *Rpt1* allele hemizygous, and thereby haplolethal, by omitting the 3’UTR from *Rpt3*. This line is characterized by 3xP3-EYFP with all remaining recoding identical to dRP. We named this design **dRPi** (**<u>d</u>**ocking site in **<u>R</u>**ibosome with **<u>P</u>**roteasome **<u>i</u>**nhibited) (**Figure 1B**).

These two docking site designs permit two broadly different gene drive types. Gene drives in dRP theoretically should permit population replacement and should be capable of driving to allelic fixation (**Figure S1A**). However these drives would experience less stringent selective pressures to guarantee homing due to tolerance of *r2* alleles in *Rpt1*, which could inhibit drive spread. Drives inserted into dRPi would however prevent r2 alleles in *Rpt1*, but would be homozygous lethal due to Proteasome knockout. We argue this does not preclude powerful replacement drive designs. In a drive design we term Recessive Lethal Replacement (RLR), a homozygous lethal gene drive could reach population fixation as heterozygotes when spread by a single sex (**Figure S1B)**. This drive would have the added benefit of simultaneous 50% population suppression in conjunction with population replacement in heterozygous form, and could thereby still meet anti-pathogen goals with inclusion of any dominant-acting antipathogen cargo [8]. Importantly both docking sites could also equally permit suppression or replacement drive designs, depending on the drive cargo, part of the elegance of these designs.

### Constructing dRP and dRPi through HACK

Direct embryonic microinjections to integrate dRP and dRPi by HDR appeared to be lethal, as we did not obtain any transformants despite repeated attempts (n=2732 embryos injected, n=245 embryos survived, 8.9% survival, compared to normal ∼25% survival). We therefore set out to harness the endogenous Interlocus Gene Conversion (IGC) phenomenon to knock-in the dRP and dRPi templates into *RpL11-Rpt1* in the germline of transgenic adults. IGC is a naturally occurring gene knockin method which templates repair not from an exogenously provided template but from distal loci. For this we developed two donor transgenes, dRP^pBdonr^ and dRPi^pBdonr^ that contain the dRP and dRPi cassettes respectively, homology arms flanking *RpL11-Rpt1*, 8 gRNAs targeting the wild type 3’ termini of *RpL11-Rpt1* coding sequences (gRNA_L-S_), and a second fluorescent selectable marker *Act5C-DsRed* outside of the Homology arms (**Figure 1C**). These donors were integrated into the genome randomly by piggyBac transgenesis and were crossed to Cas9 (VZC) [35] to generate hybrid male mosquitoes capable of IGC in the germline, (+/VZC ; +/dRP^pBdonr^) and (+/VZC ; +/dRPi^pBdonr^) genotype. The offspring larvae from these males were visibly scored for the expression of their respective *3xP3-FP* cassettes (CFP and EYFP in dRP and dRPi respectively) unlinked from *Act5c-DsRed*, indicating integration of dRP or dRPi into *RpL11-Rpt1* and dislinkage from dRP^pBdonr^ and dRPi^pBdonr^(**dRPi shown, Figure 1D**). Integration was confirmed by PCR and sequencing (**Figure 1E, 2B**). Following these experiments, we adopted the acronym Homology Assisted Crispr Knockin (HACK) to describe the phenomenon of IGC-mediated gene knockin [36,37].

We performed a brief analysis of the conditions required for HACK to aid the construction of future similar drive designs. Five dRP^pBdonr^ families with different insertion sites were assayed for their ability to template HACK of dRP into the endogenous *RpL11-Rpt1* locus (**Figure 1F**) (dRP^pBdonr^_[1]_, dRP^pBdonr^_[2]_, dRP^pBdonr^_[3]_, dRP^pBdonr^ _[4]_, dRP^pBdonr^_[5]_). Two of the families, those on the same chromosome arm as the *RpL11-Rpt1* target (Families dRP^pBdonr^_[4],_ and dRP^pBdonr^_[5]_), yielded HACK-positive larvae when screened for by *3xP3-CFP*-positive and *Act5c-DsRed*-negative fluorescence [38]. However the frequency of HACK-positive larvae was significantly higher when assayed by PCR among those who co-inherited dRP^pBdonr^, suggesting linked activity (**Figure 1G**). In these larvae, HACK of dRP into *RpL11-Rpt1* and its associated *3xp3-CFP* marker was ‘masked’ due to coinheritance with the dRP^pBdonr^ transgene containing both *3xP3-CFP* and *Actin5c-DsRed* cassettes. This suggests HACK occurred frequently in these families, but dislinkage of fluorophores through recombination or homing occurred rarely, suggesting post-Meiosis I activity. HACK was not observed visually via fluorescence from dRP^pBdonr^ donors localized to a different chromosome than the *RpL11-Rpt1* target, however PCR was not performed to identify if integration was occurring in the absence of fluorophore dislinkage. However, the dislinkage of dRP^pBdonr^ from newly integrated dRP in these lines should have allowed for identification of integrants readily via fluorescence if it was occuring. Characterization of HACK in the dRPi^pBdonr^ families was not undertaken.

In sum, HACK-based integration of both separate dRP and dRPi templates occurred from different dRP^pBdonr^ and dRPi^pBdonr^ donor loci. Taken together, these findings suggest that HACK is a reproducibly robust technology for engineering genetically intractable haplolethal loci, and most likely occurs when donor and target are linked. Furthermore these findings also demonstrate a critically important principle that homology-based repair can be tolerated in the *RpL11-Rpt1* locus when stimulated by cleavage via gRNA_L-S_.

### Characterization of dRPi and insertion of drive-like transgenes

We next set out to characterize dRPi, but chose to do so in conjunction with inserted genetic cargoes, as the binary fluorophores more easily permit identification of homozygotes at the dRPi site. Into dRPi (3xP3-EYFP fluorophore) we transgenically inserted via RMCE a drive-like transgene termed gDLT (**g**RNA-expressing **D**rive-**L**ike **T**ransgene; *3xP3-DsRed* fluorophore) (**Figure 2A, Table S2**), characterized by a non-expressing truncated Nanos promoter (**Figure 2C**) [39] upstream of Cas9, and containing expression cassettes for gRNA_L-S_ (**Figure 2A**). Integration of gDLT in dRPi was confirmed by PCR (**Figure 2B**). Crossing gDLT to dRPi enabled identification of transhomozygotes by fluorescence (gDLT/dRPi) which died prior to the first larval molt (n=28/28 died). Further analysis revealed that they experienced significant accumulation of polyubiquitin aggregates before death (**Figure 2D**), possibly due to knockout of *Rpt1* caused by the lack of replacement 3’UTR (**Figure 1A**). Importantly, gDLT represents a transgene with expression cassettes, size, and cargoes reminiscent of a functioning gene drive - including gRNAs - as it spans 1.7kB removed and 12.5kB inserted by RMCE into the docking site (**Table S2**). This suggests that the dRP and dPRPi docking sites may be able to tolerate integration of gene drives, and that the intergenic distance between *RpL11* and *Rpt1* is not of critical importance at the scales required for drive integration.

**Figure 2.**
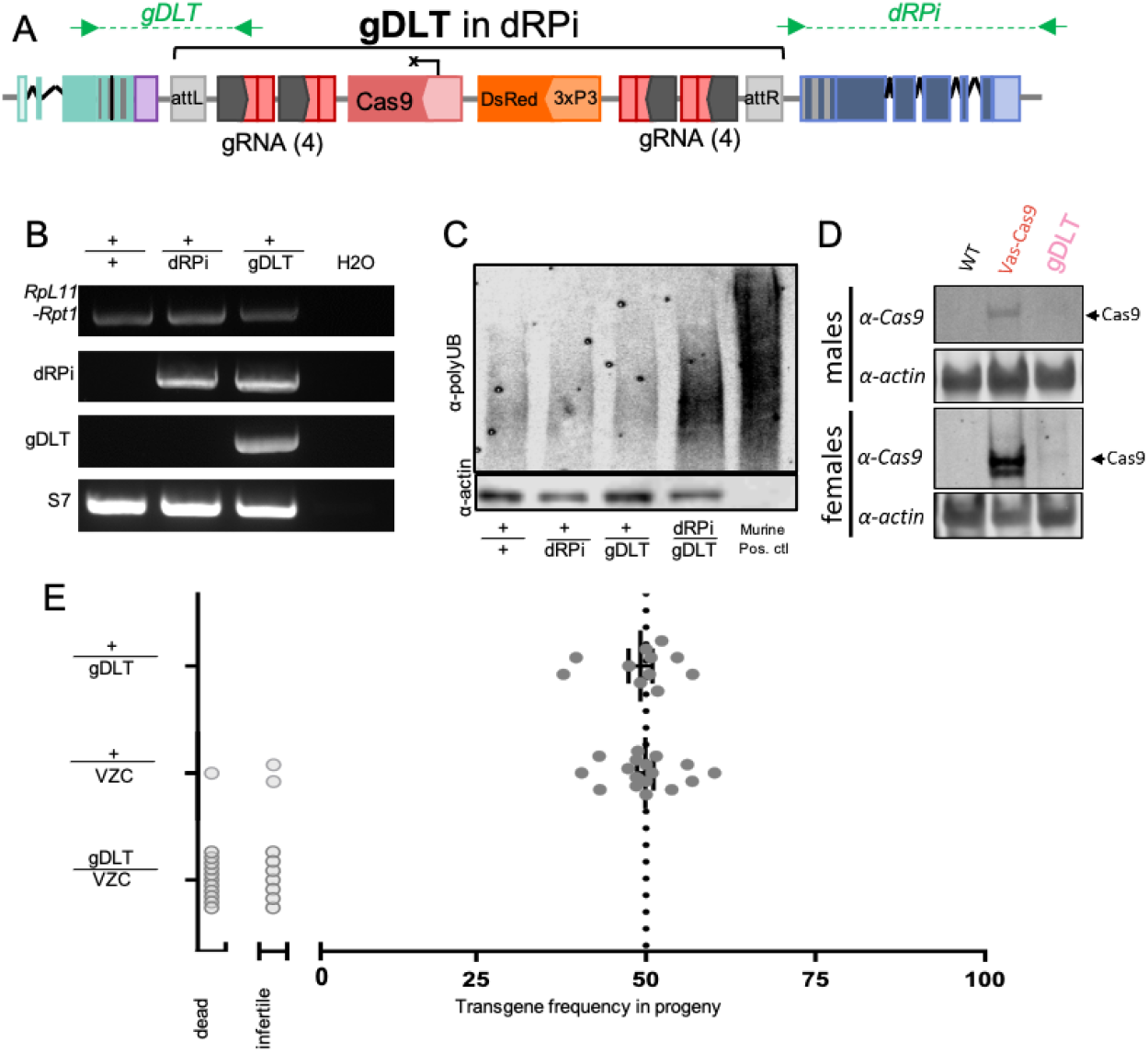
The recessive-lethal dRPi docking site, construction, characterization, validation, and testing drive phenotypes. **[A]** The transgene map of the drive-like transgene, gDLT, inserted into the dRPi docking site. **[B]** PCR validation of dRPi integration in heterozygotes, shown with wild type, dRP, dRP^pBdonr^, and water controls. *RpL11/Rpt1*: Primers specific to wild type *RpL11/Rpt1* locus spanning 1569 bp between C-terminal ends of *RpL11* and *Rpt1* removed during recoding. Primers shown in Figure 1A. *dRPi:* primers specific to integration spanning from internal sequence to genomic sequence beyond the homology arm of *Rpt1*. Predicted product size of 1922 bp for dRPi (Figure 1). *gDLT*: Primers amplify from the wild type sequence of RpL11, over the recoding and ΦC31 recombination sequence, and into the gRNA cassettes on gDLT, 1620 bp predicted size. *S7:* Control for DNA quality, 500 bp product expected. Primers not shown. **[C]** dRPi one-day-old homozygous larvae experience significant poly-ubiquitin aggregation before death compared to heterozygous and wild type siblings confirming proteasome dysfunction (SDS Nupage gel, 40 larvae per sample, 11.375 µg protein loaded each genotype, actin loading control shown). Anti-actin also shown. No band in murine positive control. **[D]** Western blot targeting Cas9 shows protein production in VZC transgenics but not in gDLT transgenics **(E)** Rescuing Cas9 function of gDLT with VZC causes infertility or death in all transheterozygous females compared to transgenic controls (p<0.0001, One Way ANOVA).

### Failure to observe homing due to lethality and sterility

We next generated multiple gene drive constructs for integration into dRPi. All constructs expressed gRNA_L-S_, and differed only in the sequences regulating Cas9, including Vasa[41], B2-tubulin[42] and two iterations of the ZPG[3] promoter. However, following injections of 7,014 dRPi and 1,012 dRP embryos, only 7 transgenics were recovered which all died as first instar larvae, suggesting insertion of gene drive transgenes into these sites may be lethal. These transgenes presumably expressed Cas9 as the promoters were identical to those previously published, while gDLT does not express Cas9 (**Figure 2C**). This suggests that the docking site may be sensitive to toxic Cas9 overexpression from inserted transgenes, possibly due to adjacent ribosomal enhancers [43].

We then assayed the drive potential of gDLT by providing Cas9 *in trans* in a split-drive-like design, crossing to the VZC *Vasa2-Cas9* line [35,44]. In a cross of transheterozygous males (+/gDLT; +/VZC) mated to wild type females, the resulting broods were completely sterile (0% fertility, n = 0/482 eggs laid, Binomial two-tailed p<0.0001), compared to control crosses of (+/VZC) sibling males which displayed normal fertility (89.9% fertility, n = 1321/1469 eggs laid, Binomial two-tailed p<0.0001). Females with the drive genotype (+/gDLT; +/VZC) experienced significant mortality 24H post-blood feeding (n = 10/15), impaired oviposition (n = 4/15), or oviposition of infertile clutches (n = 1/15)(**Figure 2E**, p<0.0001 One Way ANOVA). Taken together, these findings demonstrate significant perturbation of the reproductive biology of driving individuals consistent with significant mutagenesis of *RpL11-Rpt1*, and suggesting that CRISPR expression must be finely tuned in these designs - whether transgenically from within the docking sites or when provided *in trans* in a Split-drive-like formation - to permit fertility and observable drive.

### dRP homozygotes are Ribosome Minute mutants

During the course of experiments we discovered that dRP is homozygous inviable - a phenotype only expected of dRPi due to proteasome knockout. dRP-positive larvae reached pupation significantly more slowly (**Figure 3A**), most (dRP/dRP) individuals failed to survive eclosion (**Figure 3B**), (dRP/dRP) were significantly smaller as adults (**Figure 3C**), and females failed to develop normal ovaries following blood feeding - even in the absence of CRISPR (**Figure 3D**)[45,46]. These findings point towards (dRP/dRP) individuals being Ribosome Minute mutants with growth-defect phenotypes indicative of ribosome scarcity [47]. In concordance with this, *RpL11* mRNA levels are significantly reduced in (dRP/dRP) (**Figure 3E**). We postulate this may be a result of recoding, differential regulation caused by the *Rpt3* 3’UTR, or deletion of a necessary intergenic enhancer during docking line construction. Importantly, there were no significant differences in expression of *Rpt1* in this line (**Figure 3F**), and no observable accumulation of Poly-ubiquitin aggregates in these individuals (**Figure 3G**), indicating that the recoded Proteasome is functioning correctly as designed and the defect likely lies with *RpL11*.

**Figure 3.**
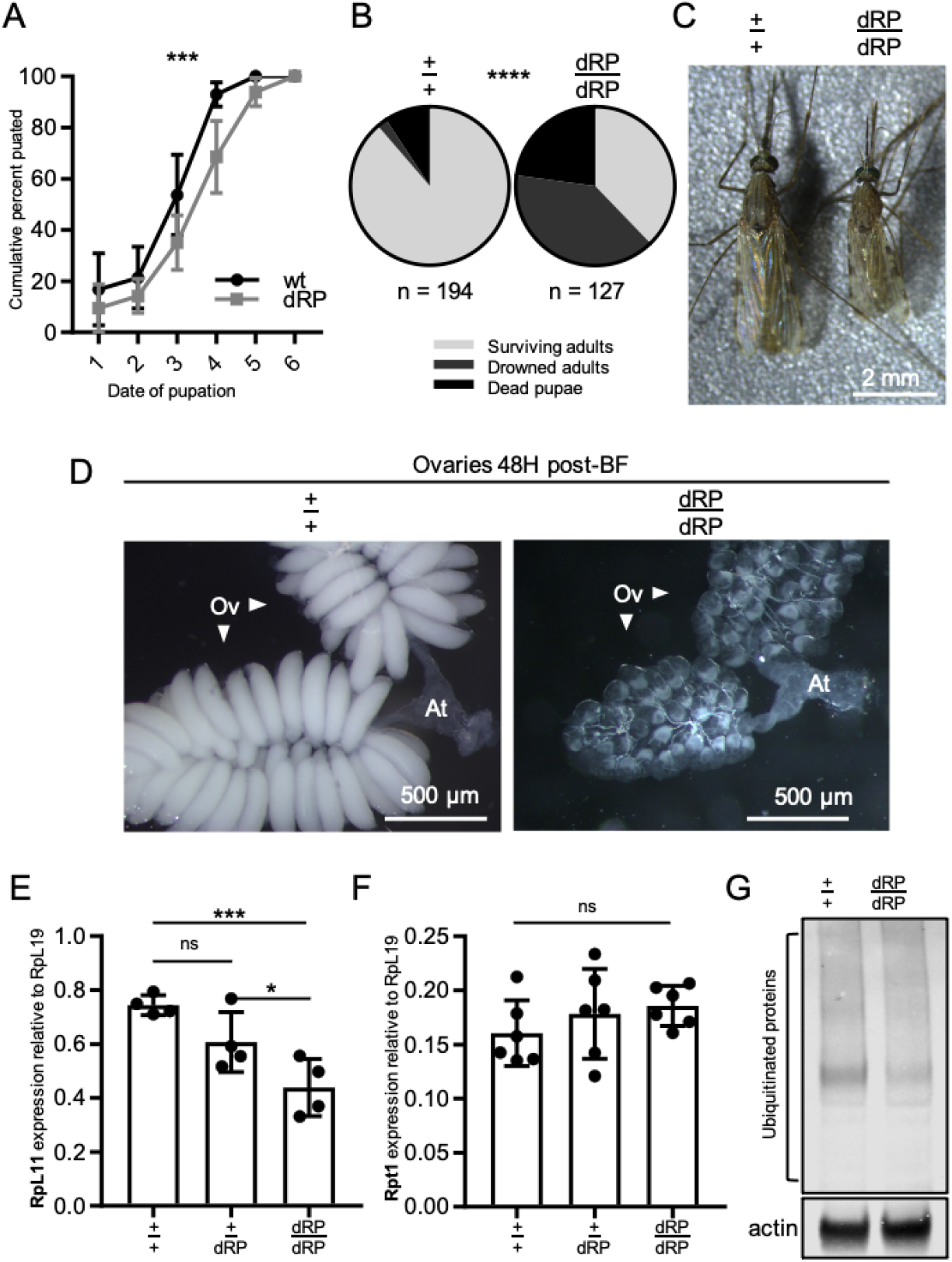
Characterization of the dRP phenotype and observation of ribosome defects. In dRP *Rpt1* proteasome function appears normal, but *RpL11* ribosome function is aberrant causing *Minute* phenotypes. **[A]** dRP-positive larvae individuals pupate later than wild type siblings. (Larval competition trays, 50 individuals per genotype, three biological replicates). Percent of total puparium formed summed each day with day 1 marking observation of first pupa. Mean and SD shown, plotted as inverse survival curve, Wilcoxon Rank; p<0.001. **[B]** Significantly fewer dRP homozygotes survive pupation than wild type siblings. Genotypes sorted as L3/L4 larvae, allowed to pupate, then scored for eclosure or mode of death within 36 h. *Surviving adults*; successfully eclosed and flew from surface of pupal dish. *Drowned adults*; fully formed adults emerged from pupal casing but drown on surface of water. *Dead pupae*; pupa dead with no movement, or individuals failed to fully emerge from pupal case before death. (Chi-squared, p<0.0001). **[C]** dRP homozygotes are smaller than wild type if they survive pupation (5-day-old female adult siblings, scale bar; 2 mm). Interestingly, some are sufficiently small that they only develop 4 legs. **[D]** dRP homozygous female ovaries fail to undergo normal oogenesis compared to wild type controls which complete vitellogenesis and develop oocytes by 48 h post-blood meal. Ov points to each of two ovarian lobes, and At denotes the atrium (equivalent to the uterus). dRP homozygotes bloodfed twice to guarantee vitellogenic surplus, controls received a single blood-feed (75x magnification, scale bar; 500 µm). **[E]** Homozygote dRP express significantly less *RpL11* mRNA transcript than heterozygous and wild type siblings (ANOVA p = 0.0035). Heterozygotes and wild type individuals have similar expression levels (one-tailed t-test, p = 0.0585), while homozygotes express significantly less than heterozygous siblings (one-tailed t-test, p = 0.0293), with highly significant 41% decrease in expression between wild type and homozygotes (one-tailed t-test, p = 0.0008). **[F]** *Rpt1* mRNA transcript levels in pupae are nonsignificantly different between dRP sibling genotypes (qRTPCR, four biological replicates per genotype, two pupae per sample, non-significant difference between groups (p = 0.3878 ANOVA, mean and SD). **[G]** There are no significant accumulations of poly-ubqiquitin aggregates in dRP homozygous pupae compared to wt siblings. (Western Blot, two pupae per sample, 40 ug per well)

## DISCUSSION

Developing gene drives that are resistance-proof may be necessary to achieve malaria control on a continental-wide scale [3]. Here we take the first steps towards development of potentially more resistance-proof HGDs in *Anopheles gambiae* by developing gene drive docking sites in a haplolethal Ribosome-Proteasome locus. We demonstrate reproducible knockin of the docking sites dRP and dRPi into the *RpL11-Rpt1* locus by HACK, and demonstrate survivability despite significant 3’ terminal recoding. In recoding, we undertake some of the first targeted whole-body engineering of endogenous ribosome genes in eukaryotes, laying the groundwork for not only gene drives, but study into the ribosome’s role in everything from ecdysone signaling, to oncogenesis to prion folding [48–50]. We discover that dRP displays a Ribosome Minute phenotype - (**Figure 3**), and characterize the proteasome knockout phenotype characteristic of dRPi (**Figure 2D**), the first such phenotypes in Anophelines to our knowledge (**Figure 2D**). We demonstrate that these docking sites are capable of RMCE insertion of a drive-like transgene, gDLT (**Figure 2A-D**), however they cannot currently harbor fully functioning gene drives likely due to a combination of Cas9 toxicity and Ribosome-depletion phenotypes (**Figure 3)**.

Though beyond the scope of this work, different strategies could be used to demonstrate gene drive of dRP or dRPi. Fine-tuning Cas9 expression using insulators, codon-deoptimization, induction systems, or nickases could be employed, in addition to novel promoter expression systems [5]. Assaying for homing as a split-drive design could enable characterization of necessary components in isolation, and providing an additional transgenic copy of recoded *RpL11* could alleviate Minute phenotypes. In designing these systems such that only successful homing permits the cell to survive, it is clear that more finely-tuned Cas9 is required to enable drive while not causing lethality or toxicity of surrounding germline tissue. Beyond traditional homing gene drives, these docking sites could also enable construction of CLvR or HomR systems with one of the recoded genes providing the necessary gene rescue gene product for function [51,52].

HACK was used to generate these lines, and may provide a valuable tool for engineering genetically intractable loci for future drive systems. Two different donor sites were used successfully to generate dRP and separately dRPi was generated using a unique donor line, suggesting HACK is recapitulatable so long as the donor is on the same chromosome as the recipient locus. This technology promises to be useful for creating drives in intractable loci in the future.

In all, these lines represent a significant advancement in the development of haplolethal-targeting gene drive designs, and present important avenues for future study. They represent the most advanced haplolethal-targeting drive designs in Anophelines to date, and with future optimization may form the basis of powerful gene drive designs. Despite the absence of drive, our work still presents a major advancement in knowledge in the phenotypes and construction of haplolethal-targeting gene drive designs, providing important insights for the field of vector control and similar drive designs being developed in other metazoans.

## METHODS

### gRNA design

8 gRNAs were designed and selected based on their *in silico* cleavage prediction (https://zlab.bio/guide-design-resources). All gRNAs were designed in 4 pairs, two pairs each targeting the 3’ terminus of each gene, and were designed in such a way to enable cleavage with a Cas9 nickase to generate overhangs to further bias for HDR. gRNAs L and M were synthesized as a tethered pair with alternative scaffold sequences from S. pyogenes and S. mutans. Their sequence can be found in **Table S3**. gRNAs N and O were synthesized as a tethered pair with alternative scaffold sequences from *S. Pyogenes* and *S. agalactiae*. Their sequence can be found in **Table S3**. gRNAs R and S were synthesized as a tethered pair with alternative scaffold sequences from *S. Pyogenes* and *S. agalactiae*. Their sequence can be found in **Table S3**. gRNAs P and Q were synthesized as a tethered pair with alternative scaffold sequences from *S. mutans* and *S. pyogenes*. Their sequence can be found in **Table S3**.

### Recoding

Recoding was undertaken manually using the codon optimization table found in Volohonski et al [34]. All codons were recoded to the next-most common codon, giving preference to the most common codons. Codons with <20% frequency were omitted. Surrogate 3’ UTRs from *RpL32* and *Rpt3* were manually selected to replace the endogenous *RpL11* and *Rpt1* 3’UTRs. They were chosen based on subunit stoichiometry in respective protein complexes and a general understanding of protein function. Full sequences can be found in **Table S1**.

### Cloning and transgenesis

Cloning of dRP^pBdonr^, dRPi^pBdonr^, and gDLT was undertaken using standard molecular biology protocols including Golden Gate and Gibson. These plasmids were established transgenically following standard molecular biology protocols and were used to generate dRP and dRPi lines respectively (see “**Interlocus gene conversion crosses to establish dRP and dRPi”** below). Due to the Pandemic, these plasmids are no longer available. Therefore the full plasmid sequence with feature annotation is available in **Supplementary Table 1 and 2**. The difference between dRP^pBdonr^ and dRPi^pBdonr^is omission of the *Rpt3* 3’UTR. Additional gRNA target sequences were included in the plasmid to amplify cleavage by a second generation of gene drives at this site not otherwise discussed. gRNA cleavage sites for a second generation gene drive are noted as gRNAL’,M’,N’,O’,P’,Q’,R’, S’ in **Supplementary Table 1**. Second generation gene drives are however not further discussed in this work. Vasa-Cas9, VZC, transgene was reported previously [35]. The transgene gDLT sequence is provided and annotated in **Supplementary Table 2**.

### Inverse PCR on dRP^pBdonr^ to determine integration loci

Inverse PCR was essentially as described in [53]. In essence, 1-3uL of genomic DNA was digested with TaqI enzyme for 4h. This was then circularized with ligase in a 100uL reaction. The sample was concentrated by precipitation via Sodium Acetate then was resuspended in 10uL water. 1uL of this preparation was used as a template for PCR. PCR was carried out with the primers 1114H.S3 and 1114H.S4, (5’ CTGTGCATTTAGGACATCTCAGTC 3’) and (5’ GACGGATTCGCGCTATTTAGAAAG 3’) respectively, the latter of which amplifies outwards beyond the piggyBac terminal repeat and into adjacent genomic sequences. PCR amplicons were gel extracted, cloned into pJET (Thermo Scientific, Cat. No. / ID: K1231), and individually sequenced.

### Interlocus gene conversion crosses to establish dRP and dRPi

F0 VZC males and dRP^pBdonr^, dRPi^pBdonr^ females were crossed together en masse (approximately 50-100 individuals each genotype). They produced F1 hybrid {+/VZC; +/dRP^pBdonr^} or {+/VZC ; +/dRPi^pBdonr^} offspring. Among these F1 offspring, males were outcrossed to WT in mass (approximately 50-100 individuals each genotype), and the F2 offspring were screened for integration of dRP or dRPi by fluorescence. Fluorescence was visible as 3xP3-CFP or 3xP3-EYFP in the absence of Actin5c-DsRed respectively for dRP and dRPi.

### Western Blots

Western blots were carried out essentially as described in [54]. For Cas9 Westerns 7 5-10 day old male lower abdomen or female ovaries were prepared, and for Poly-ubiquitin Westerns, 30 1 day old larvae were genotyped by fluorescence and and prepared in Np-40 cell lysis buffer system (ThermoFisher Scientific) supplemented with 1mM PMSF in DMSO and 40µ Protease Inhibitor. Western blots to detect Cas9 were carried out with α-Cas9 [Cell Signaling Technologies ®(Cas9 XP ®, Rabbit mAB # 19526, 1:1000 dilution)]. Westerns targeting poly-ubiquitin failed to identify an anopheline positive control, therefore a murine tissue culture sample with known proteasome defects was used as a positive control which does not stain for actin (1µg/µL C2C12 cell line treated with Bortezomib). Membranes were treated with 6M GuHCl, 20mM Tris pH 7.5, 1mM PMSF and 5mM βME for 30 min at 4° to release Poly-ubiquitinated tails from aggregates [55], before blocking and antibody incubation with 1:500 solution of the Fk2 Mono- and polyubiquitinated antibody (Enzo Life Sciences, BML-PW150-0025).

### Genotyping PCR

Genotyping PCRs were undertaken on single 4-8 day old adults following standard Molecular Biological protocols. PCR for the presence of the WT *RpL11-Rpt1* allele was undertaken with primers als561 and als287, [5’ CGGGCATGTTTGCGATTC 3’] and [5’ CTGACCAAGGAGGACGCG 3’] respectively, spanning the 3’ terminal coding region of wild type *RpL11* and *Rpt1* for an amplicon of 1577 bp. PCR for the presence of the integrated docking site, dRP or dRPi, was amplified with primers als328 and als306, [5’CACACCTTTACATATCGCTCGC 3’], [5’ GTACCTTCCAGGTCGTAGTCTTG 3’], spanning the *Rpt1* 5’ UTR outside of the homology arms and the length of the *Rpt1* coding sequence. The amplicon is 2,193 bp in dRP, and 1,995 bp in dRPi due to the presence or absence of the surrogate *Rpt3* 3’UTR respectively. The dRP^pBdonr^ was PCR amplified with primers als287 and als478, [5’ CTGACCAAGGAGGACGCG 3’] and [5’ GTGATGGTCACGGTGCTTTTAC 3’], respectively, amplifying the gRNAs and Actin5c promoter, for a 916 bp fragment. Control S7 PCRs were amplified with FC107 and FC108, [5’ GGCGATCATCATCTACGTGC 3’] and [5’ GTAGCTGCTGCAAACTTCGG 3’], targeting the wild type ribosomal S7 subunit. The gDLT primers als 291 [5’GCGGTCAACAAGGTGACAAGG 3’] and als342 [5’ CTGCGTCGCGACAACTTCTC 3’] amplify from the wild type sequence of *RpL11*, over the recoding and ΦC31 recombination sequence, and into the gRNA cassettes on gDLT, 1620 bp predicted size.

### qRT-PCR

Samples for quantitative RT-PCR were two 3 day old adults, one male one female, diluted tenfold and quantified in triplicate using standard curves. PCRs were run in Fast SYBR Green Master Mix (Thermo-Fisher) on a Step One Plus thermocycler (Applied Biosystems).

qRT-PCR amplification of *RpL11* was undertaken with als1024 and als1025, [5’ TAAGGTAGCCACAATGCCAGC 3’] and [5’GCGCATCACGTTCTTCGACT 3’] respectively, targeting the first exon-exon junction and overlapping the start codon, an unmodified sequence shared by both the recoded and wt alleles of *RpL11*. qRT-PCR amplification of Rpt1 was undertaken with als1018 and als1019, [5’ AGGTGAACGAACTGACGGG 3’] and [5’ CCTGAAGCGGCTGCTCATT 3’] targeting exon 3 of *Rpt1*, a shared sequence in both recoded and wt *Rpt1*. Quantities were normalized against the ribosomal protein RpL19 using previously described primers[56].

### Assaying for inheritance bias

Females carrying either the +/gDLT, +/VZC, or +/gDLT;+/VZC phenotype were mated *ad libitum* to wild type males for 3 days. They were then bloodfed and isolated into individual oviposition cups to lay their eggs. The resulting larval broods were then scored for the number of fluorescent larvae over the total number of larvae. They were then plotted as a percentage of transgenic offspring over total. Females which died or laid wholly infertile broods are plotted in open dots at left.

### Delayed pupation assay

Broods resulting from an intercross of +/dRP x +/dRP parents were raised with approximate larval density of 200 larvae per tray. Within any given brood, the number of transgenic or wild type individuals pupating each day was scored. Transgenic individuals included both +/dRP and dRP/dRP hetero- and homozygotes. The data is plotted as an inverse survival curve with day 6 corresponding to the last date of pupation and a percentage of 100%

### Pupal mortality assay

dRP/dRP and WT sibling individuals were identified and separated as L4 larvae by fluorescence intensity. They were allowed to pupate and were checked over the course of 2 consecutive days. The number of successfully eclosed, dead, and drowned adults were counted.

### Ovarian dissection

Female dRP/dRP and WT were genotyped as L4 larvae, then allowed to eclose into adults. At 3 days old they were provided a blood meal to induce oogenesis. 48h later the reproductive tract was dissected and imaged under a Leica Stereomicroscope.

## Data availability

Complete sequence maps and plasmids are available in Table S1 and S2. Transgenic lines dRP and dRPi are upon request to O.S.A.

## Acknowledgments

We thank Emily Lund, Reema Apte, James Pai, Sansa Chen, Martha Chow, Michelle Bui, Julika Job, and Simon Joseph for helping with mosquito husbandry.

This research was funded by the Howard Hughes Medical Institute/Bill and Melinda Gates Foundation Grant OPP1158190 to F.C; by the National Institutes of Health (NIH) (award number R01 AI104956 to F.C.; and by an F31 AI120480-02 to A.S. F.C. is funded by the Howard Hughes Medical Institute (HHMI) as an HHMI investigator. Defense Advanced Research Projects Agency under the Safe Gene program to K.E. and G.C.; Burroughs Welcome Fund IRSA 1016432, and NIH R00-DK102669-04 to K.E. This work was supported by funding from NIH awards (R01AI151004, RO1AI148300, RO1AI175152) awarded to O.S.A. The findings and conclusions within this publication are those of the authors and do not necessarily reflect positions or policies of the HHMI or the NIH. The funders had no role in the study design, in data collection, analysis or interpretation, in the decision to publish, or the preparation of the manuscriptFigures were created using www.BioRender.com.

## Author Contributions

A.L.S conceptualized and designed experiments, performed molecular analysis and genetic experiments, analyzes and compiled the data, wrote the manuscript draft and contributed to final manuscript editing; E.A.M., S.S., and E.M. performed molecular analyses, and genetic experiments; O.S.A provided mentorship, provided husbandry support, and edited the manuscript. G.M.C. provided significant intellectual contributions and edited the manuscript. F.C provided mentorship, significant intellectual contributions, and edited the manuscript. K.M.E designed experiments, provided significant intellectual contributions, and edited the manuscript. All authors approve the final manuscript.

## Ethical conduct of research

All animals were handled in accordance with the Guide for the Care and Use of Laboratory Animals as recommended by the National Institutes of Health and approved by the UCSD Institutional Animal Care and Use Committee (IACUC, Animal Use Protocol #S17187) and UCSD Biological Use Authorization (BUA #R2401).

## Disclosures

O.S.A is a founder of Agragene, Inc. and Synvect, Inc. with equity interest. The terms of this arrangement have been reviewed and approved by the University of California, San Diego in accordance with its conflict of interest policies. G.M.C. has the following patents related to this work: WO2015006290A1 (“Multiplex RNA-guided genome engineering”) and WO2016089866A1 (“RNA-guided systems for in vivo gene editing”). A complete list of G.M.C’s conflict of interest can be found at arep.med.harvard.edu/gmc/tech.html. K.M.E. and A.L.S have pending patents related to the described work: WO2015105928A1 (“RNA-guided gene drives”) and K.M.E. has an additional patent WO2015006294A2 (“Orthogonal Cas9 proteins for RNA-guided gene regulation and editing”). All other authors declare no competing interests

**Supplementary Figure 1.**
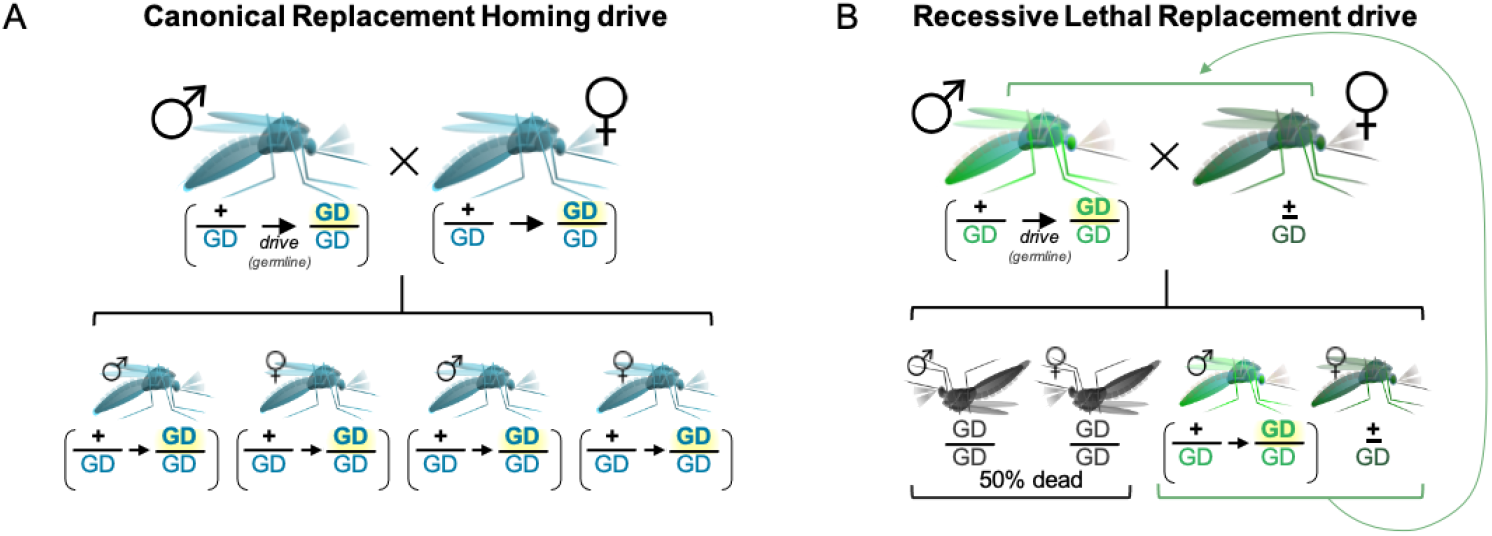
**[A]** Canonical population replacement gene drives home in the germline of both sexes and permit survival of all individuals to facilitate population fixation of the drive allele and any associated cargo. dRP was originally designed to enable this type of population replacement drive. However this drive is not possible with the lines discussed herein due to unexpected Ribosome Minute phenotypes **[B]** Late stages of recessive-lethal replacement (RLR) drive heterozygotes intercross, resulting in death of half the offspring and heterozygote dominant fixation of drive individuals. Drive occurs in the germline of a single sex (males here, bright green) contributing (up to) 100% drive-positive chromosomes to the next generation, while heterozygote survival is permitted by the contribution of a wild type chromosome from the non-driving sex (females here, dark green), which recapitulates the parental cross *ad infinitum* (curved green arrow). Concurrent death of homozyogtes (dark grey) each generation causes simultaneous population suppression by half.

